# Biomarker Localization, Analysis, Visualization, Extraction, and Registration (BLAzER) Workflow for Research and Clinical Brain PET Applications

**DOI:** 10.1101/608323

**Authors:** Fabio Raman, Sameera Grandhi, Charles F. Murchison, Richard E. Kennedy, Susan Landau, Erik D. Roberson, Jonathan McConathy, Alzheimer’s Disease Neuroimaging Initiative

## Abstract

**Objective:** There is a need for tools enabling efficient evaluation of amyloid- and tau-PET images suited for both clinical and research settings. The purpose of this study was to assess and validate a semi-automated imaging workflow, called Biomarker Localization, Analysis, Visualization, Extraction, and Registration (BLAzER). We tested BLAzER using two different segmentation platforms, FreeSurfer (FS) and Neuroreader (NR), for regional brain PET quantification in images from participants in the Alzheimer’s Disease Neuroimaging Initiative (ADNI) dataset.

**Methods:** 127 amyloid-PET and 55 tau-PET studies along with corresponding volumetric MRI were obtained from ADNI. The BLAzER workflow utilizes segmentation of MR images by FS or NR, then visualizes and quantifies regional brain PET data using FDA-cleared software (MIM), enabling quality control to ensure optimal registration and detect segmentation errors.

**Results:** BLAzER analysis required only ∼5 min plus segmentation time. BLAzER using FS segmentation showed strong agreement with ADNI for global amyloid-PET standardized uptake value ratios (SUVRs) (r = 0.9922, p < 0.001) and regional tau-PET SUVRs across all Braak staging regions (r > 0.97, p < 0.001) with high inter-operator reproducibility for both (ICC > 0.97) and nearly identical dichotomization as amyloid-positive or -negative (2 discrepant cases out of 127). Comparing FS vs. NR segmentation with BLAzER, the global SUVRs were strongly correlated for global amyloid-PET (r = 0.9841, p < 0.001), but were systematically higher (4% on average) with NR, likely due to more inclusion of white matter, which has high florbetapir binding.

**Conclusions:** BLAzER provides an efficient workflow for regional brain PET quantification. FDA-cleared components and the ability to visualize registration reduce barriers between research and clinical applications.

## INTRODUCTION

Positron emission tomography (PET) neuroimaging applications have increased in both research and clinical setting in recent years. PET provides the ability to study functional and molecular processes in the brain *in vivo*, allowing exploration of an array of normal and pathological states, including neurodegenerative disorders, psychiatric conditions, and neurooncology. Quantitation of PET data is well-established for research applications but less so in routine clinical settings. Recent research demonstrates the utility of quantification to supplement visual assessment for clinical PET, especially in Alzheimer’s disease (AD) using the glucose analogue 2-deoxy-2-[^18^F]fluoro-D-glucose (FDG) and amyloid PET tracers [1-4]. Quantification of targeted brain regions of interest (ROIs) [5, 6] is enabled using masks generated by automated magnetic resonance imaging (MRI) segmentation algorithms, such as the widely used reference standard in research, FreeSurfer (FS) [7-9], or the FDA-cleared and ISO-certified Neuroreader (NR) [10-13]. By registering the segmentation mask with the PET scan, ROIs can be defined for PET measurements without laborious and potentially imprecise or biased manual delineation methods [6, 14]. Barriers to the use of this approach to quantitation of brain PET, particularly for clinical applications, include computationally intensive software, time-consuming workflows, the need for real-time visualization of the primary and processed data for quality control, and limited availability of FDA-cleared software suitable for this purpose.

Accuracy and precision are key features of effective image analysis methods to ensure that the extracted biomarker data can provide reliable *in vivo* information of the physiological state or disease process of interest. However, efficiency, ease of use, and availability are also essential features for widespread and routine implementation of quantitative brain PET. Many current techniques require lengthy post-processing time [2, 15] and/or heavy computational workload [16, 17]. Incorrect segmentation of the brain MRI data and misregistration between PET and segmented brain regions are potential sources of error when extracting of regional PET data based on ROIs defined through fully automated segmentation [15]. Additionally, although qualitative and quantitative assessment can be performed separately, the ability to easily visualize images in real-time at different states of processing and registration can provide an important degree of quality control and allow for user input to correct problematic studies. Many of the current fully automated processing workflows that involve only inputs and outputs to optimize speed, can be liable to unidentified errors because there is no opportunity to visualize the processed images [18, 19]. Finally, other barriers to routine implementation of quantitative brain PET include the lack of versatility in existing tools to analyze images acquired or processed by different platforms, radiotracers, and segmentation algorithms. Radiotracers commonly used for brain PET analysis in AD include [^11^C]Pittsburgh compound B (PiB) [20], [^18^F]florbetapir [21, 22], [^18^F]florbetaben [23], and [^18^F]flutemetamol [24] for amyloid-β (Aβ), [^18^F]flortaucipir for tau [25-28], and [^18^F]FDG [29, 30] for glucose metabolism. Multicenter clinical trials and longitudinal studies can benefit from a versatile workflow to standardize PET quantification [31].

Although a number of effective research workflows exist for regional brain PET analysis that combine one or more of the previously discussed features [8, 19, 32-35], quantification still has not been widely implemented in routine clinical brain PET for AD, which relies primarily on visual assessment and components validated for routine clinical use. FDA clearance and ISO-certification are key features of not only NR, but also MIM, software that provides image visualization and quantification capabilities. Combining features of such tools could help lower the barriers between clinical and research workflows. However, to the best of our knowledge, no workflow exists that is efficient and easy for physicians and technologists to use, automatically segments ROIs in a customizable fashion, consists of FDA-cleared components, and allows for both qualitative and quantitative assessment for brain PET data in a clinical environment.

We present here a novel brain Biomarker Localization, Analysis, Extraction, and Registration (BLAzER) workflow for analysis of PET based on segmented brain MRI. We demonstrate that BLAzER works well for both global and regional PET quantification of brain amyloid- and tau-PET, respectively, in Alzheimer’s Disease Neuroimaging Initiative (ADNI) participants spanning normal cognition, mild cognitive impairment (MCI), and AD dementia. Although this report focuses on AD biomarkers, BLAzER can be applied to any PET neuroimaging study that utilizes MR-based segmentation for defining ROIs. Additionally, we compare two different inputs for segmentation, FS and NR, which could be used with BLAzER for research and clinical applications, respectively.

## METHODS

### Study Population

Images were downloaded from the Alzheimer’s Disease Neuroimaging Initiative (ADNI) database (http://www.loni.ucla.edu/ADNI/). ADNI was launched in 2003 as a public-private partnership, led by principal investigator Michael W. Weiner, MD. The primary goal of ADNI has been to test whether serial magnetic resonance imaging (MRI), PET, other biological markers, and clinical and neuropsychological assessment can be combined to measure the progression of mild cognitive impairment (MCI) and early AD. All studies were approved by local Institutional Review Board at each institution through ADNI. Additional information is available at the ADNI website (www.adni-info.org).

127 amyloid-PET and 55 tau-PET studies were selected from ADNI for a total of 178 unique subjects (4 overlap). Subjects were selected to represent the spectrum of ADNI participants in terms of cognitive status (cognitively normal (CN), mild cognitive impairment (MCI), and Alzheimer’s disease (AD)), and age (cohorts aged 55-59, 60–69, 70–79, and ≥80). Subject selection was performed prior to beginning image review or analysis and was based on the preceding demographic criteria alone to capture the broad range of neuropathology available in ADNI (Table 1). Selection was blinded to the actual quality of the scans and output of the data, and image analysis was performed blind to subjects’ cognitive status or age group. No exclusions were made at time of selection or subsequently during the analysis (i.e. all of the initially selected studies were included in the final analysis).

**Table 1.** Cohort demographics. Subjects in each of the three groups, cognitively normal (CN), mild cognitive impairment (MCI), and Alzheimer’s disease (AD), were stratified into four age groups. Number of total subjects (N), broken down by sex with number of males (M). Apolipoprotein E-ε4 (ApoE4) carriers represented as either homozygotes (ε4/ε4) or heterozygotes (ε4/ε*). All other results are shown as mean ± standard deviation (SD). * indicates incomplete data in ADNI.

|  | CN | MCI | AD |
| --- | --- | --- | --- |
| **Amyloid-PET with florbetapir** |  |  |  |
| N (Male, %) | 37 (13M, 13%) | 48 (21M, 43.8%) | 42 (27M, 64.3%) |
| ApoE4 carriers (homo, hetero, total %) | 11 (0, 11, 31.6%) | 30 (5, 25, 62.5%) | 27 (10, 17, 65.9%) |
| Age in years (mean ± SD) | 76 (±8.8) | 70.7 (±10.6) | 71.9 (±9.9) |
| MMSE (mean ± SD) | 28.5 (±1.8) | 23.8 (±6.1) | 22.95 (±2.1) |
| ADAS11 (mean ± SD) | 5.9 (±4.1) | 16.9 (±12.9) | 18.9 (±6.5) |
| ADAS13 (mean ± SD) | 9.4 (±6.3) | 25.1 (±16.9) | 29.1 (±8.1) |
| ADNI Global SUVR (mean ± SD) | 1.14 (±0.13) | 1.22 (±0.1) | 1.3 (±0.2) |
| **Tau-PET with flortaucipir** |  |  |  |
| N (Male, %) | 19 (7M, 36.8%) | 19 (15M, 78.9%) | 16 (10M, 62.5%) |
| ApoE4 carriers (homo, hetero, total %) | 4 (0, 4, 22.2%)* | 7 (3, 4, 36.8%) | 7 (3, 4, 50.0%)* |
| Age in years (mean ± SD) | 77.74 (±6.9) | 77.89 (±6.1) | 79.19 (±9.3) |
| MMSE (mean ± SD) | 29.47 (±0.6) | 27.47 (±2.7) | 21.25 (±4.4) |
| ADAS11 (mean ± SD) | 7.42 (±3.9) | 11.74 (±5.5) | 22.06 (±8.1) |
| ADAS13 (mean ± SD) | 10.63 (±5.8) | 17.58 (±8.0) | 32.31 (±11.2) |

### Image sets used for analysis

All subjects in both the amyloid-PET and tau-PET cohorts had a volumetric brain MRI scan performed within 1 year of the PET study. MRI studies were performed using standard T1-weighted sagittal 3D MPRAGE (magnetization prepared rapid gradient echo) sequences acquired from various 3T scanners with a 1.25 × 1.25-mm in-plane spatial resolution and 1.2-mm slice thickness with 256 x 256 voxel resolution according to ADNI specifications.

Amyloid- and tau-PET studies were acquired with [^18^F]florbetapir (also called AV-45) and [^18^F]flortaucipir (also called AV-1451), respectively. PET images were obtained on different scanners of varying resolutions, each with its platform-specific acquisition protocol. Therefore, as part of the ADNI protocol, all raw PET images undergo pre-processing for quality control and standardization purposes at the University of Michigan [36]. In summary, 4 × 5 min dynamic image frames, acquired 50 to 70 minutes post-injection, were co-registered to the first extracted frame of the raw image file. Then, the 6 five-minute frames were averaged to form a static PET image and reoriented into a standard 160 × 160 × 96 voxel image grid with 1.5-mm cubic voxels. Finally, each image set was filtered with a scanner-specific filter function to produce images of uniform isotropic resolution of 8 mm FWHM, which is the approximate resolution of the lowest resolution scanners used in ADNI. Only the fully pre-processed, standardized, co-registered, and averaged PET images were used for this study. Further details on the ADNI acquisition protocol are available on the website (http://adni.loni.usc.edu/methods).

### Image Analysis

#### MRI Segmentation

Volumetric MRI images were segmented by either FreeSurfer v6.0.0 (Boston, MA) or Neuroreader (Brainreader, Horsens, Denmark). FS, an open-source software, has been validated as a tool to measure brain volumes in various neurological diseases when compared to either manual delineation [6] or other algorithms [14]. FS uses a complex algorithm with a series of normalization and motion-correction approaches prior to even starting its intensity-based approach to parcellate the brain into not only the larger cortical structures, but also subregions to allow targeted measurement of specific ROIs [8, 9]. FS segmentation produces detailed ROIs, which allow for spatial delineation of brain regions beyond the main cortical and subcortical brain structures. Customization of brain regions is particularly useful in analysis of tau-PET studies in Alzheimer’s disease where pathology follows characteristic Braak staging regions [37].

NR’s separation of gray matter (GM), white matter (WM), and cerebral spinal fluid (CSF) uses an intensity-based approach along with a template, similar to other available approaches [38, 39] such as the widely used Automatic Anatomic Labeling (AAL) template [19]. Similar to AAL, NR is highly efficient, with segmentation of an individual case processed in approximately 15 minutes compared to 8 to 12 hours for FS. However, segmented brains from NR provide larger cortical and subcortical brain regions compared to FS, limiting customization beyond anatomical boundaries defined by their atlas.

FS studies were loaded on a supercomputer environment (Cheaha, Birmingham, AL) in order to run all 184 volumetric MRI input files simultaneously through parallel computing. Parcellated, segmented brain images were converted from MGZ to DICOM format using 3DSlicer v4.6 (Boston, MA). Alternatively, NR automatically produced segmented brains in DICOM format for the same MPRAGE scans. Each of the segmentation output files along with source volumetric MRI and pre-processed PET scans were then visualized and quantified using an automated workflow on a multi-modal imaging software – MIM v6.6.13 (MIM Software Inc., Cleveland, OH).

#### PET Analysis

MIM is a commercial, FDA-cleared software package designed to help researchers and clinicians quantitatively and qualitatively process multi-modal imaging data. Through a graphical user interface (GUI), MIM allows the user to design customized, automated workflows for processing cases without need for advanced computational skills. Workflows were developed that could use segmentation data from FS or NR as inputs for ROI definition on the PET data sets. For BLAzER, we selected the segmented brain dataset, brain PET, and volumetric brain MRI images for analysis, then ran a user-defined and customizable workflow that registered the images, defined ROIs within the original volumetric MR and PET based on the segmentation, and delivered the quantification in an array of potential outputs while allowing for visual inspection. Workflows were generated and customized to an individual segmentation method and PET radiotracer through a GUI interface where steps in the protocol were simply selected from a drop-down list similar to Porcupine [40], such that users were not limited by their computational skillset to analyze large, multi-modal image datasets. Once the original protocol had been generated, anatomical tags for brain regions from another segmentation method or radiotracer were incorporated by replacing with the new identifiers with minimal user input, allowing clinicians and researchers alike to implement their customized analyses through BLAzER with minimal training.

In summary, in the BLAzER workflow, we 1) segmented the MRI to generate a 3D brain mask using either FS or NR; 2) transferred the segmentation to MIM and selected corresponding volumetric MRI and PET to analyze; 3) automatically co-registered segmented brain mask to volumetric MRI; 4) utilized the segmented brain to delineate the various brain regions based on the pixel intensities for each region defined by FS or NR; 5) performed quality control through visualization and corrected registration, if necessary; and 6) fused PET scan to MRI/brain mask template to extract, visualize, and quantify data (Fig. 1). The workflow suspended the automated process before transferring the contours to allow the user to verify accurate image registration. The original MPRAGE was used as a quality control measure to assure that the rigid registration method used to align the images had been performed properly and the ROIs are correct. If needed, the user could manually correct the registration and/or ROIs at this step. This review step was employed for all cases but is optional and could be omitted to provide a more automated process.

**Figure 1.**
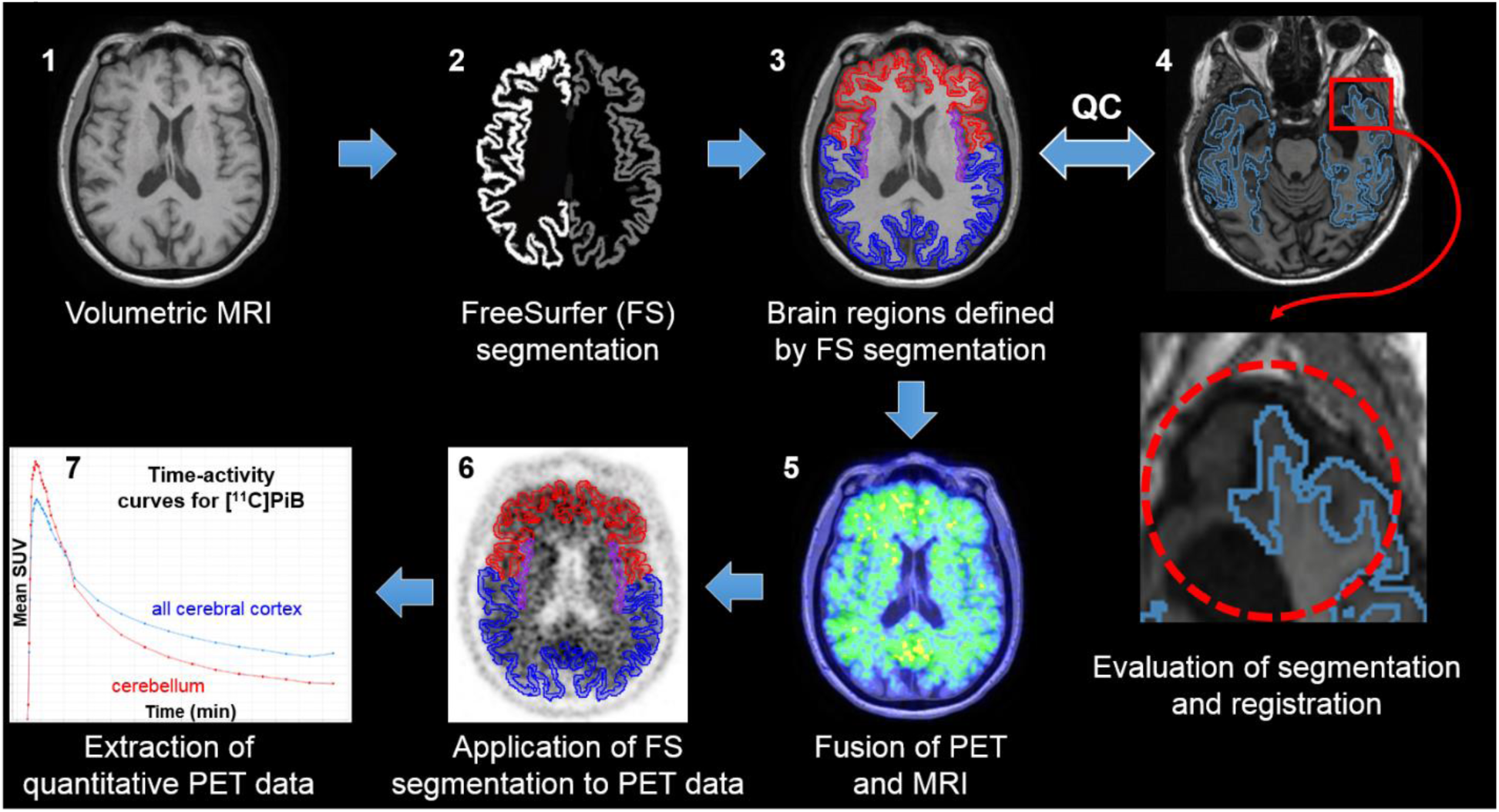
Schematic diagram of key steps in the semi-automated BLAzER workflow. The 3D volumetric MRI (1) is used to generate the Freesurfer (FS) segmentation (2) in DICOM format. Brain regions defined by FS are generated in MIM and applied to the original MRI (3). Quality control is performed (4) to ensure both registration and segmentation are correct. In an example from a different research subject, a cortical region in the anteromedial left temporal lobe is not included in the FS-defined segmentation (red circle showing left temporal cortex not outlined by blue line). Once any errors are corrected, the PET and MRI data are fused (5). The FS-defined regions are transferred to the PET data set (6) which are then used to extract static or dynamic regional PET data (7).

#### Anatomical Definitions of Brain Regions

In BLAzER, all SUVRs extracted are automatically weighted by the volumes of the individual subregions comprising each ROI. Amyloid-PET ROIs were normalized to the entire cerebellum to extract SUVRs for all regions that comprised the cerebral cortex: frontal lobe, temporal lobe, parietal lobe, and cingulate (BLAzER-FS, as defined by ADNI) or frontal, temporal, parietal, and occipital lobes (BLAzER-NR, as defined by Neuroreader). BLAzER-FS matched ADNI-defined cortical regions exactly while BLAzER-NR included the entire cortex. For head-to-head comparisons between BLAzER and ADNI, BLAzER-FS delineation was based on ADNI-defined brain subregions. However, for comparisons between BLAzER-FS and BLAzER-NR, BLAzER-FS anatomical regions were modified to match the NR-based regions of entire cortex. We denote this distinction as BLAzER-FS*. Specific FreeSurfer subregions are detailed below as well as listed in Supplementary Table 1.

Specifically, BLAzER-FS subregions were defined as follows: caudal middle frontal, lateral orbitofrontal, medial orbitofrontal, pars opercularis, pars orbitalis, pars triangularis, rostral middle frontal, superior frontal, and frontal pole (frontal); caudal anterior cingulate, isthmus cingulate, posterior cingulate, and rostral anterior cingulate (cingulate); inferior parietal, precuneus, superior parietal, and supramarginal (parietal); middle temporal and superior temporal (temporal). In contrast, BLAzER-FS* regions were defined as follows: caudal middle frontal, lateral orbitofrontal, medial orbitofrontal, pars opercularis, pars orbitalis, pars triangularis, rostral middle frontal, superior frontal, frontal pole, paracentral gyrus, precentral gyrus, caudal anterior cingulate, posterior cingulate, and rostral anterior cingulate, and insula (frontal); inferior parietal, precuneus, superior parietal, supramarginal, postcentral gyrus, and isthmus cingulate (parietal); bankssts, entorhinal, fusiform, inferior temporal, middle temporal, parahippocampal, superior temporal, temporal pole, and transverse temporal (temporal); cuneus, lateral occipital, lingual, and pericalcarine (occipital).

Tau-PET SUVRs were calculated similarly to the amyloid-PET defined ROIs with two main differences. Tau-PET was normalized to the cerebellar gray matter instead of the entire cerebellum [37] and ADNI-defined ROIs for BLAzER-FS workflow followed the pathologic Braak staging regions [41]: entorhinal cortex (Braak 1); hippocampus (Braak 2); parahippocampal, fusiform, lingual, and amygdala (Braak 3); middle temporal, caudal anterior cingulate, rostral anterior cingulate, posterior cingulate, isthmus cingulate, insula, inferior temporal, and temporal pole (Braak 4); superior frontal, lateral orbitofrontal, medial orbitofrontal, frontal pole, caudal middle frontal, rostral middle frontal, pars opercularis, pars orbitalis, pars triangularis, caudate, putamen, lateral occipital, parietal supramarginal, inferior parietal, superior temporal, pallidum, superior parietal, precuneus, superior temporal sulcus, nucleus accumbens, and transverse temporal (Braak 5); pericalcarine, postcentral, cuneus, precentral, and paracentral (Braak 6). As NR’s anatomical atlas only includes major cortical and subcortical regions, we defined BLAzER-NR and BLAzER-FS* tau-PET regions the same as for amyloid-PET: frontal, temporal, parietal, and occipital cortices.

The published ADNI data was used as the reference standard. The reported values in the ADNI data set were based on brain segmentations performed using FS v5.3.0. SUV and SUVR data was extracted using SPM5 (version) after co-registeristration of PET and MR data. Key differences between BLAzER and the ADNI method are summarized in Table 2.

**Table 2.** Key differences in post-processing methods summarized between BLAzER and the ADNI reference standard.

| **BLAzER-FS** | **ADNI** |
| --- | --- |
| **Segmentation using FS v6.0.0** | **Segmentation using FS v5.3.0** |
| **One MPAGE scan utilized for segmentation** | **Two independent MPAGE scans utilized for segmentation** |
| **Registration performed by MIM** | **Registration performed by SPM** |
| **Visualization step for quality control to ensure adequate registration and manual/automated editing if necessary** | **Registration fully automated** |
| **Dichotomized data both using ADNI's 1.11 cutoff as well as performing regression to determine corresponding "BLAzER" units** | **Derived cutoff autopsy studies of Clark et al. (34). Converted their 1.10 cutoff to 1.11 "ADNI" units based on regression analysis** |

### Statistical Analysis

Statistical analyses were performed using MedCalc v17.7.2 (MedCalc Software, Mariakerke, Belgium) and Matlab vR2016b (MathWorks, Natick, MA) to compare the BLAzER method with the reference standard from ADNI. SUVRs and volumes were reported as mean ± SD. Statistical significance was defined as *p*<0.002 to account for multiple comparisons. Univariate regression was used to validate measurements between the two methods using Pearson’s correlation coefficient (poor agreement=0; slight=0.01-0.20; fair=0.21-0.40; moderate=0.41-0.60; good=0.61-0.80, and excellent=0.81-1.00 agreement). Similar comparisons were conducted using Bland Altman to determine limits of agreement and percent differences between the two analysis tools. Intra- and inter-observer variability and agreement were evaluated for global cortical SUVR using the intraclass correlation coefficient (ICC) in a two-way random model (ICC<0.40 = poor; ICC≥0.40 to 0.75 = fair to good; ICC>0.75 = excellent agreement). Finally, amyloid-PET cases were dichotomized into positive or negative to determine the ability of BLAzER to properly classify cases based on SUVR cutoffs. Dichotomization was performed by 1) directly applying ADNI’s autopsy-derived 1.11 cutoff [34, 42] and 2) deriving BLAzER-specific cutoff by using a linear regression to convert “ADNI units” to BLAzER units based on the slope and y-intercept.

## RESULTS

### Efficiency, quality control, and reproducibility

The BLAzER workflow enabled rapid image analysis. The most time-consuming step was segmentation. NR was considerably faster than FS on a per-case basis (10–20 min/case vs. 8–12 hours/case, respectively), yet slower than FS for total processing time for the 182 subject cohort (45.5 hours vs. 12 hours, respectively) due to the ability to run FS cases in parallel on a supercomputer environment. After segmentation, FS cases required ∼2 min for manual DICOM conversion whereas NR cases were returned in DICOM format. Once the segementation was obtained, the remainder of BLAzER processing took about 5 min per case. Thus, a case could be fully processed, from completion of image acquisition to full regional quantification, in as little as 20 min (when using NR segmentation).

Although the workflow could operate with full automation, the ability to visualize registration provided for quality control and avoiding registration errors. We routinely ran MIM’s “Run Rigid Assisted Alignment” tool serially to fix minor errors until it provided no further adjustment, which took only a few seconds (Fig. 1). Only 1 of the 182 scans required manual correction of registration, which was a case with severe brain atrophy. This highlights the usefulness of the visualization step, however, as the incorrect registration would have been missed in a fully automated workflow. We also examined inter-rater reliability by having two independent operators process each scan. Global florbetapir- and regional flortaucipir-PET measurements showed excellent reproducibility between two users across all brain regions (ICC > 0.97, Table 3).

**Table 3.** Inter-subject reproducibility of BLAzER. Interclass Coefficient (ICC) values show excellent reproducibility between two different operators.

| Region | florbetapir-PET | Volumetric MR |
| --- | --- | --- |
| Frontal | 0.9975 | 0.9914 |
| Parietal | 0.9915 | 0.9961 |
| Temporal | 0.997 | 0.9944 |
| Cingulate | 0.9961 | 0.9713 |
| Global | 0.9974 | 0.9986 |
| Region | flortaucipir-PET |  |
| Braak 1 | 0.9847 |  |
| Braak 2 | 0.9833 |  |
| Braak 3 | 0.9963 |  |
| Braak 4 | 0.9993 |  |
| Braak 5 | 0.9924 |  |
| Braak 6 | 0.9679 |  |

### MRI Volumetric Measurements

As BLAzER’s regional PET extraction depends on accurate MRI-based anatomic segmentation and registration, we first compared BLAzER measurements of regional MRI volumes to the ADNI reference standard. Global cortical volume measured by the BLAzER workflow was highly correlated to that measured by ADNI (r = 0.9749, p < 0.001, Fig. 2A) with small systematic difference (1.61%) and tight 95% confidence interval (CI) (Fig. 2B). Regional comparisons across the frontal, cingulate, parietal, and temporal lobes for CN, MCI, and AD subjects showed similar results (r > 0.92, p < 0.0001, Supplementary Table 2).

**Figure 2.**
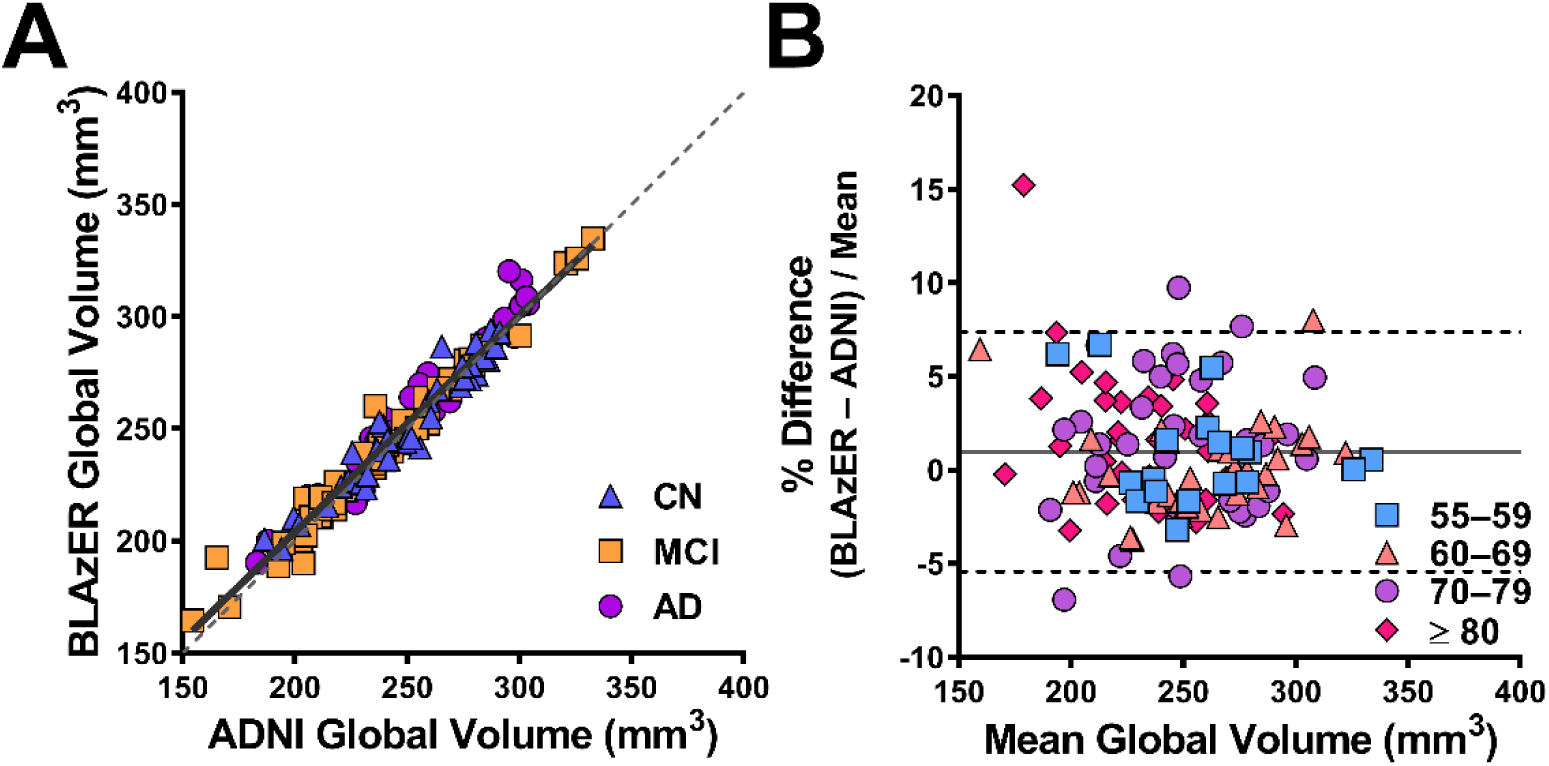
Comparison of MRI cortical volumes determined by BLAzER-FS vs. ADNI. Global cerebral cortical volumes are based on frontal, temporal, parietal, and cingulate cortices as defined in ADNI. A) Univariate Pearson correlation, with regression line (r = 0.97, solid black), identity line (dotted), and cases coded by group. B) Bland-Altman plots with mean percent difference (0.98%, solid line) and 95% confidence intervals ([–5.43 to 7.38], dotted lines), coded by age cohort.

### Amyloid-PET SUVR and Dichotomization

BLAzER showed strong agreement with the ADNI reference standard when measuring diffuse cortical binding of amyloid-PET. Global SUVRs were similar between BLAzER and ADNI (r = 0.9922, p < 0.0001, Fig. 3A). Additionally, the slope of linear regression and y-intercept (y = 1.0012x + 0.01816, R^2^ = 0.9844, Table 4) showed a near one-to-one correspondence between BLAzER and ADNI. There was a slight systematic difference in global SUVR, which was on average 1.6% higher with BLAzER than in the ADNI dataset (Fig. 3B). This may be to do a systematic reduction in the reference region mean, which was observed in the cerebellum. Regional comparisons across the frontal, cingulate, parietal, and temporal lobes for CN, MCI, and AD subjects showed similar results (r > 0.94, p < 0.0001, Supplementary Table 3).

**Table 4.** Dichotomization results for amyloid-PET for BLAzER-FS vs. ADNI. Number discrepant represent the total number of individuals whose dichotomous classification differed between BLAzER-FS and ADNI out of 127 total cases. Dichotomization performed by (*top row*) applying ADNI’s unadjusted autopsy-derived cutoff of 1.11 and (*bottom row*) translating this cutoff by performing a regression against the ADNI data (y = mx + b).

|  | **Number**<br>**Discrepant** | **Cutoff** | **m(x)** | **b** |
| --- | --- | --- | --- | --- |
| **BLAzER-FS using ADNI cutoff** | 4/127 | 1.11 | - | - |
| **BLAzER-FS using regression-**<br>**adjusted cutoff** | 2/127 | 1.13 | 1.0012 | 0.01816 |

**Figure 3.**
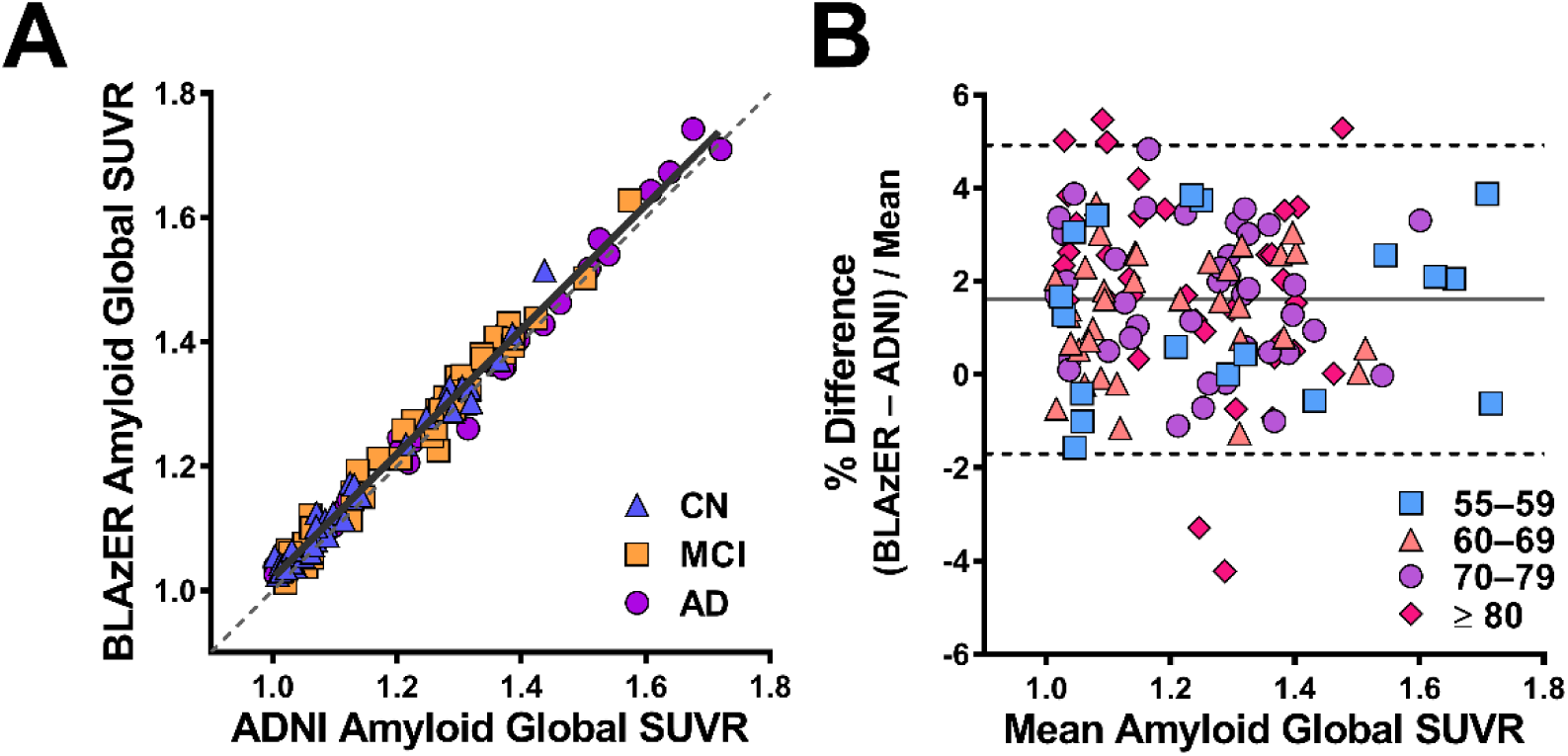
Comparison of florbetapir-PET global cerebral cortical SUVR determined by BLAzER-FS vs. ADNI. A) Univariate Pearson correlation with regression line (r = 0.99, solid black), identity line (dotted), and cases coded by group. B) Bland-Altman plots with mean percent difference (1.61%, solid line) and 95% confidence intervals ([–1.70% to 4.92%], dotted lines), coded by age cohort.

We also the examined the dichotomous classification of individuals as amyloid-positive vs. –negative based on SUVR cutoffs. Because BLAzER SUVRs were slightly higher than ADNI on average, a few more of the 127 subjects were classified as amyloid-positive when BLAzER SUVRs were dichotomized using the ADNI cutoff of 1.11 with all 4 of these cases lying near the cutoff (BLAzER vs. ADNI range: [1.113 to 1.125] vs. [1.060 to 1.098], respectively, Table 4). However, translating 1.11 “ADNI units” into 1.13 “BLAzER units” by linear regression (Equations 1 and 2) fixed the dichotomization of these 4 cases but 2 different cases were discrepant out of the 127 total at this cutoff. These 2 cases had SURVs of 1.113 vs. 1.125 and 1.113 vs. 1.115 for BLAzER-FS vs. ADNI, respectively.

1. 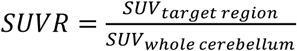

2.*SUVR* _*BLAzER*_*= SUVR* _*ADNI*_ * *slope* _*BLAzER*_ + *intercept* _*BLAzER*_

### Regional brain tau-PET comparison

BLAzER showed strong agreement with the ADNI reference standard when looking at regional binding of tau-PET with flortaucipir. Despite slightly lower correlation coefficients and wider 95% confidence intervals than with amyloid-PET (as expected because of the smaller regions defined for analysis), BLAzER correlated strongly with the ADNI reference standard across all six Braak stage regions (r > 0.97, p < 0.0001, Fig. 4). Further demonstrating regional accuracy, BLAzER showed tight 95% CI (<10%) and small differences (–1.5 to 3.6%) across all regions, (Fig. 5), and all cognitive statuses (r > 0.91, p < 0.0001, Supplementary Table 3).

**Figure 4.**
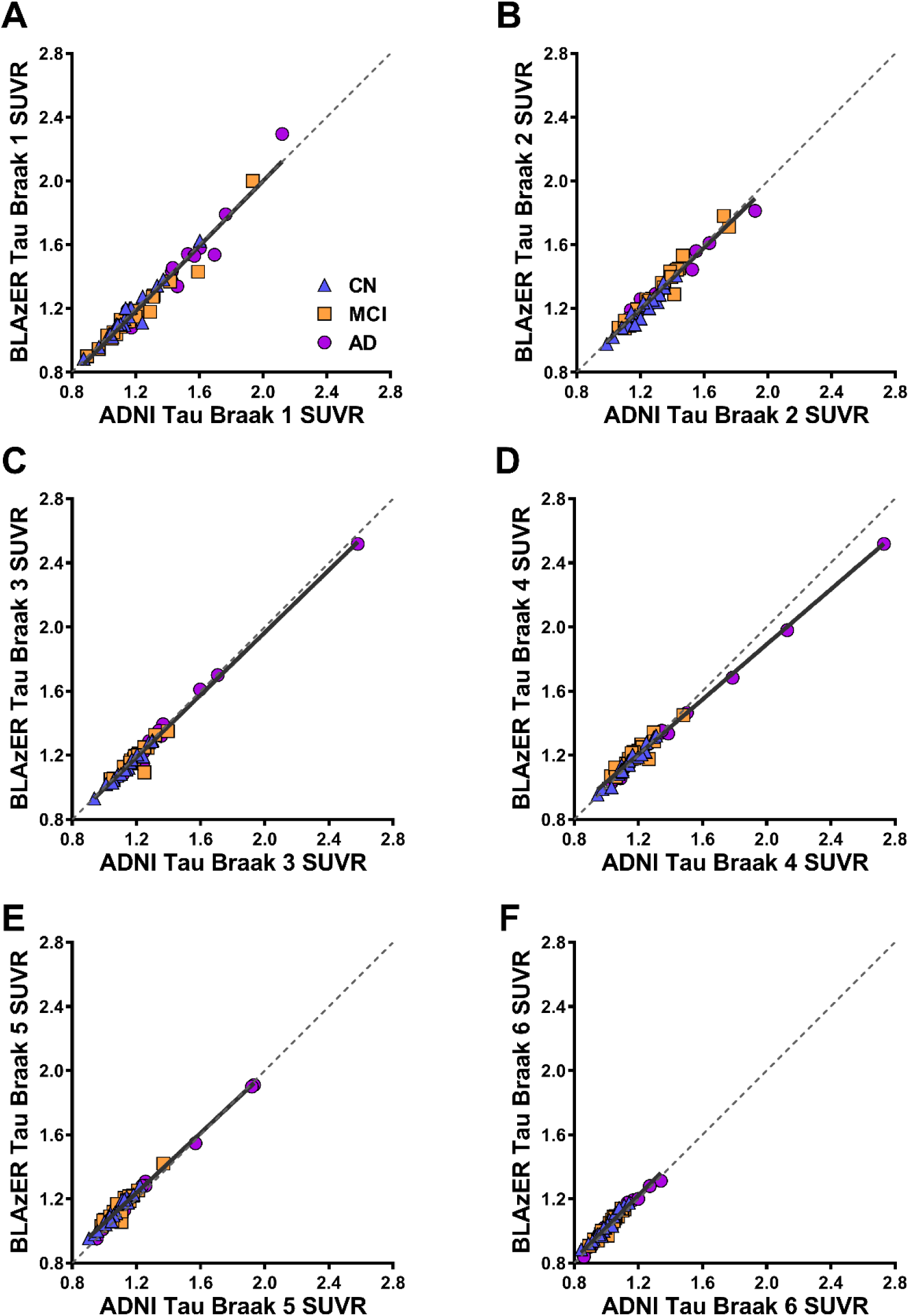
Comparison of flortaucipir-PET measured by BLAzER-FS vs. ADNI across the regions representing the pathological Braak stages. (A) through (F) represent the regions for Braak stages 1 thru 6, respectively. See Methods for details on which anatomic regions were included in each stage. Univariate Pearson correlation with regression line (solid black), identity line (dotted), and cases coded by group.

**Figure 5.**
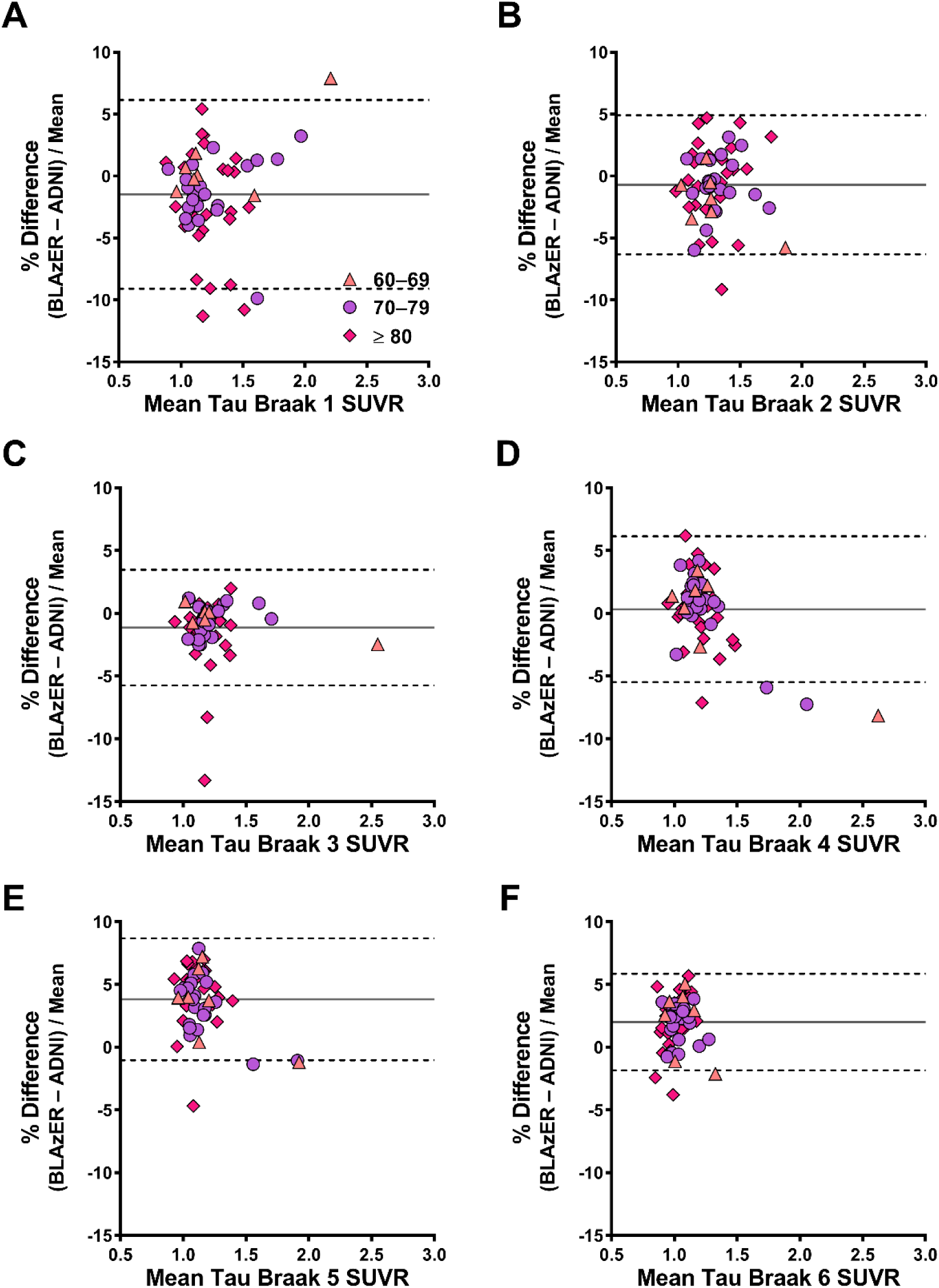
Comparison of flortaucipir-PET measured by BLAzER-FS vs. ADNI across the regions representing the pathological Braak stages. (A) through (F) represent the regions for Braak stages 1 thru 6, respectively (see Methods for details on which anatomic regions were included in each stage), Bland Altman plots with mean percent difference (solid line) and 95% confidence intervals (dotted lines), coded by age cohort.

### Amyloid and tau PET quantitation based on NR segmentation

Next we compared the performance of a different segmentation platform, comparing BLAzER-FS with BLAzER-NR. To enable apples-to-apples comparison, we first adjusted FS’s anatomical subregions to match NR’s composite regions (FS*, see methods). BLAzER-NR highly correlated with BLAzER-FS* at a global level for amyloid-PET (r = 0.9841, p < 0.001, Fig. 6A). However, BLAzER-NR values had higher SUVR than BLAzER-FS* as shown by the systematic difference (4.0%) and shifted 95% CI (–0.7% to 8.7%, Fig. 6B). This is likely due to BLAzER-NR segmentation including more PET signal from white matter, which has high florbetapir binding, due to a slightly thicker definition of cortical regions with NR compared to FS. Dichotomization results paralleled these findings (Table 5). Directly applying ADNI’s cutoff of 1.11 classified led to 12 out of 127 cases being discrepant between BLAzER-NR and BLAzER-FS* (range of SUVRs: [1.092 to 1.180] vs. [1.049 to 1.111], respectively). After applying regression analysis for both BLAzER-NR to ADNI (y = 0.9845x + 0.0.05658, R^2^ = 0.9587) and BLAzER-FS to ADNI (y = 0.8947 × + 0.1158, R^2^ = 0.9482), the new SUVR cutoff obtained was 1.15 for BLAzER-NR while BLAzER-FS* remained unchanged. The adjusted cutoff reduced the number of discrepant cases to 7 out of 127 (NR vs. FS range: [1.0923 to 1.180] vs. [1.088 to 1.120], respectively). This discrepancy reflects the differences between the segmentation provided by NR and FS which in turn affects SUVRs.

**Table 5.**
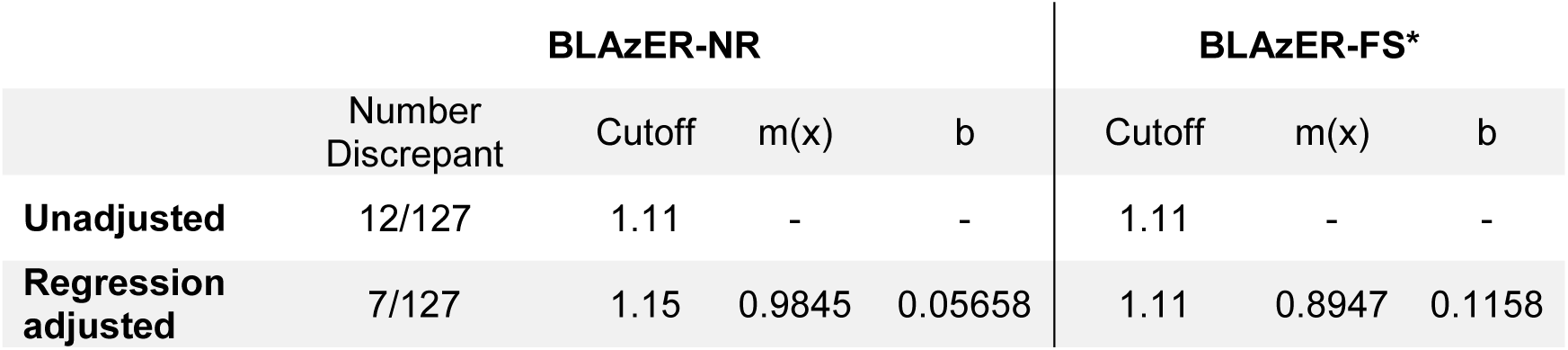
Dichotomization results for Neuroreader (BLAzER-NR) vs. FreeSurfer (BLAzER-FS*). Number discrepant represent the total number of individuals whose dichotomous classification differed between BLAzER-NR and BLAzER-FS* out of 127 total cases. Dichotomization performed by (*top row*) applying ADNI’s unadjusted autopsy-derived cutoff of 1.11 and (*bottom row*) translating this cutoff by performing a regression against the ADNI data (y = mx + b). BLAzER-FS* indicates that the BLAzER workflow used FS segmentation with brain region ROIs matched as closely as possible to those defined by NR segmentation.

**Figure 6.**
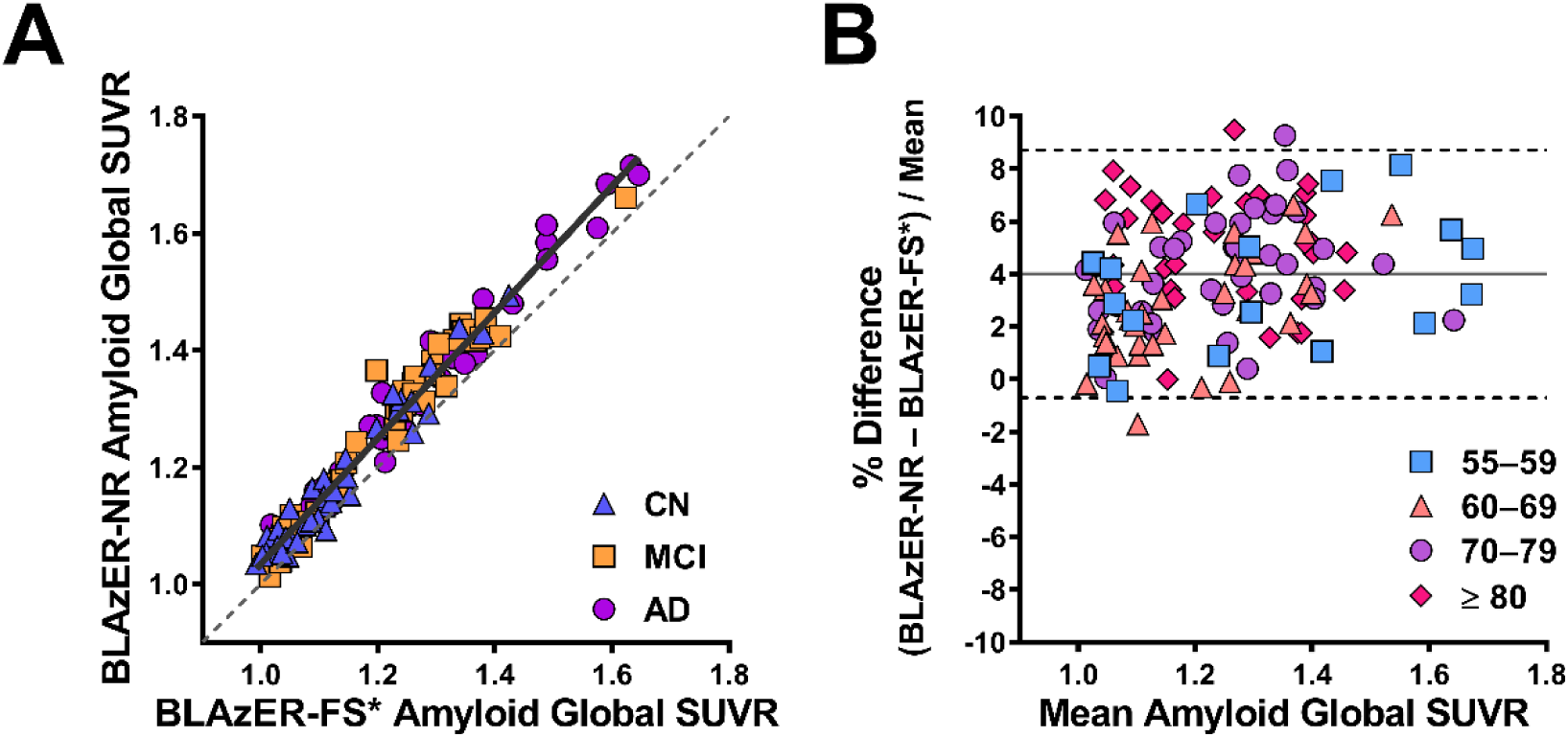
Comparison of global amyloid PET SUVR for Neuroreader (BLAzER-NR) vs. FreeSurfer (BLAzER-FS*). A) Univariate Pearson correlation with regression line (r = 0.98, solid black), identity line (dotted), and cases coded by group. B) Bland-Altman plots with mean percent difference (4.01%, solid line) and 95% confidence intervals ([–0.70% to 8.72%], dotted lines), coded by age cohort.

Regional analysis for both amyloid- and tau-PET across different cognitive statuses and brain regions revealed excellent correlations between BLAzER-NR and BLAzER-FS* (r > 0.85, p < 0.0001, Supplementary Table 4) although not as high as the correlations between BLAzER-FS and the ANDI reference values, again reflecting differences in the segmentation between the NR and FS platforms.

## DISCUSSION

Measurement of the regional brain distribution of PET tracers is a cornerstone of data analysis in molecular neuroimaging and is growing in importance for clinical applications, particularly in the evaluation of patients with cognitive impairment. A number of imaging processing workflows exist that perform quantification of regional brain PET data [8, 19, 32-35]. However, there are few if any analysis tools that perform well for both clinical and research applications, utilize FDA-cleared components, define ROIs and extract regional PET data automatically, provide both quantitative and real-time visual assessment, and work quickly and efficiently without need for advanced coding skills. Our evaluation of BLAzER with validation through comparison to the ADNI database and head-to-head comparison of two segmentation methods demonstrates an approach that addresses this unmet need.

Quantitative brain PET data from BLAzER correlated very closely with the results in ADNI for florbetapir- and flortaucipir-PET analyzed using FreeSurfer-defined regional volumes. We also demonstrated that the regional brain volumes provided by FS and NR were in close agreement with ROIs defined through BLAzER with less than 2% error between the original volumes and those output by BLAzER. BLAzER and ADNI produced nearly identical dichotomous classification of amyloid-PET participants as positive or negative, especially when using regression analysis to account for systematic differences. It is important to acknowledge that our analysis was completed before the most recent ADNI update, which includes a new partial volume corrected set of values based on modified reference regions. Thus, any partial volume effect (PVE), or inaccuracies that result from low PET spatial resolution [43], affected both BLAzER and our reference ADNI set similarly. In addition to PVE, off-target-binding in non-specific regions can be problematic for accurate PET quantification. Recent evidence has shown non-specific flortaucipir uptake in the basal ganglia [44, 45]. In the most recent ADNI updates, the caudate and putamen have been removed from the Braak 5 staging regions for flortaucipir. However, we kept our results consistent by comparing BLAzER to the original dataset that included the caudate and putamen. In future work, BLAzER-FS regions can be modified to account for PVE and off-target-binding if desired.

We observed slightly larger systematic differences and CIs for flortaucipir-PET compared to florbetapir-PET. As pathological cortical tau distribution is more regionally localized and restricted than pathological cortical amyloid which tends to be diffuse throughout the cerebral cortex [28], differences in segmentation and image alignment are expected to have a greater effect on PET measures of regional brain tau compared to regional brain amyloid.

Potential sources of the small differences between BLAzER and the ADNI data set include different FS versions, source volumetric MR images, and registration methods. For our study, FS v6.0.0 was used while v5.3.0 was used for the ADNI data set, which could affect anatomical volume measurements and brain segmentation which in turn would affect ROI definition for PET quantitation [46]. Additionally, the ADNI dataset used two volumetric MR scan input files for FS to produce the segmented brain while we chose to use only one volumetric brain MR to reflect our standard research and clinical workflows. Within ADNI, each subject undergoes two volumetric MR scans during a single scanning session with some subject datasets having slightly different imaging parameters. In order to align the PET and MR images, ADNI utilized SPM to register the images automatically while we used MIM. BLAzER utilizes a rigid-registration algorithm that allows only translation and rotation. Additionally, we utilize a three-step system of registration where the segmentation is first registered with its original, template volumetric MR and then with the PET after the rigid-registration quality control-check. We believe that this visual assessment and option for manual correction minimizes sources of error that can arise from fully automated registration methods (Fig. 1) [2]. Additionally, we believe that our results are generalizable across the ADNI population due to our selection of individuals that represent the range of cognitive statuses and age cohorts from ADNI.

Our implementation of BLAzER for research studies utilizes a high-throughput, supercomputing cluster for segmentation (by FS) along with a commercially available multimodality image viewing and analysis software program (MIM) with customized workflows to use and process the segmentation data. This workflow can be adapted to other segmentation algorithms by customizing the visualization software workflow with minimal change in the user experience. This workflow allows the user to register and process brain segmentation maps with brain PET and brain MRI in a single viewing environment for quality control, manual correction if needed, data extraction, and real-time visual assessment. Utilizing parallel computing strategies with FS allows segmentation of over 100 studies in approximately 12 hours which is suitable for use with clinical trials and large imaging datasets. We chose MIM due to its efficiency and relative ease of use compared to existing tools that are either dependent on relatively heavy computational skills [16, 17] or require significant image processing input [2, 15]. MIM’s GUI-interface is most similar to Porcupine [40], enabling analysis of large, multi-modal image datasets by users without a sophisticated computational skillset. Similar to other existing workflows [2, 47-49], BLAzER allows for automatic delineation of PET ROIs. However, an important advantage of BLAzER is that it can accept brain segmentation data sets from multiple sources, including FDA-cleared algorithms such as NR. This potential integration of a completely FDA-cleared workflow helps overcome barriers between clinical and research applications of BLAzER. While the present work focuses on the use of this workflow for amyloid- and tau-PET, it is just as applicable for the analysis of other brain PET imaging studies. This flexibility combined with visualization software familiar to physicians and technologists facilitates implementation of quantitative PET neuroimaging in routine clinical practice with a wide range of segmentation algorithms and PET tracers. As BLAzER can accommodate any DICOM dataset, other neuroimaging modalities, such as SPECT for Parkinson’s disease or functional MRI and diffusion tensor imaging for schizophrenia, could be processed along with PET neuroimaging biomarkers of neuroinflammation [50-52] by adapting the BLAzER workflow. Although not used in this data study, the BLAzER workflow can be used for analysis of dynamic PET data sets.

Another major strength of the BLAzER workflow is the ability to use any segmentation map in DICOM format for defining PET ROIs and extracting PET measurements. However, in contrast to our inter-workflow comparison between BLAzER-FS and ADNI, our intra-workflow comparison between BLAzER-FS* and BLAzER-NR demonstrated subtle differences between the segmentation methods, yet still showed excellent correlations for both global and regional comparisons. Our results showed that NR provides slightly higher SUVRs when compared to FS across large cortical structures, where dichotomous classification revealed 7 discrepant individuals out of 127 total cases even after regression adjustment. These findings, which contrast with the nearly identical dichotomization between BLAzER-FS and ADNI comparison, emphasize the effect that differences in segmentation can have on amyloid dichotomization. Despite our best efforts to match BLAzER-NR’s cortical anatomical boundaries with rearrangement of BLAzER-FS* regions, BLAzER-NR and BLAzER-FS* have slightly different anatomical definitions of brain subregions across cortical structures not only from each other but also from ADNI (unlike BLAzER-FS), which noticeably affects correlation and dichotomization. Additionally, the methods utilize different segmentation algorithms in separating gray from white matter, leading to slightly thicker cortical ROIs with NR than with FS. As florbetapir binds non-specifically to white matter even in cognitively normal individuals [53], we believe that a greater contribution of PET signal from white matter in BLAzER-NR contributes to the systematically larger SUVRs when compared to BLAzER-FS*. Users can choose segmentation software that best fits their clinical and/or research needs. Based on our observed results, the selection of the appropriate segmentation input depends on the application. For rapid analysis of large major cortical or subcortical structures in a clinical setting, NR and similar algorithms might be preferred. In contrast, FS provides more versatility for brain segmentation due its ability to provide smaller ROIs compromised of multiple subregions which may be more appropriate for research and analysis of PET tracers with spatially-restricted distributions.

In conclusion, BLAzER is a streamlined image processing workflow for efficient registration, visualization, and extraction of brain PET data. We successfully validated the accuracy and reproducibility of BLAzER using ADNI as the reference standard for amyloid-PET and tau-PET analyses. We also showed how two different segmentation inputs, NR and FS, can be used with BLAzER and that the cortical ROIs extracted by BLAzER remain consistent with the original MR volumes. This versatile workflow can reduce barriers to quantitative brain PET and MR analysis for a wide range of research and clinical applications.

## Availability of data and materials

Data used in the preparation of this article were obtained from the ADNI database (<u>adni.loni.usc.edu</u>), which is easily available for download from the Laboratory of Neuroimaging (LONI) website to the research public.

## Author Contributions

FR, EDR, and JM designed the research studies. FR and SG performed image analysis. FR, SG, CM, RK, SL, EDR, and JM interpreted data. CM, RK, SL, EDR, and JM provided methodology expertise. ADNI provided imaging studies. FR, EDR, and JM wrote the manuscript. All authors reviewed, edited, and approved the final version of the manuscript.

---

**Table S1.**
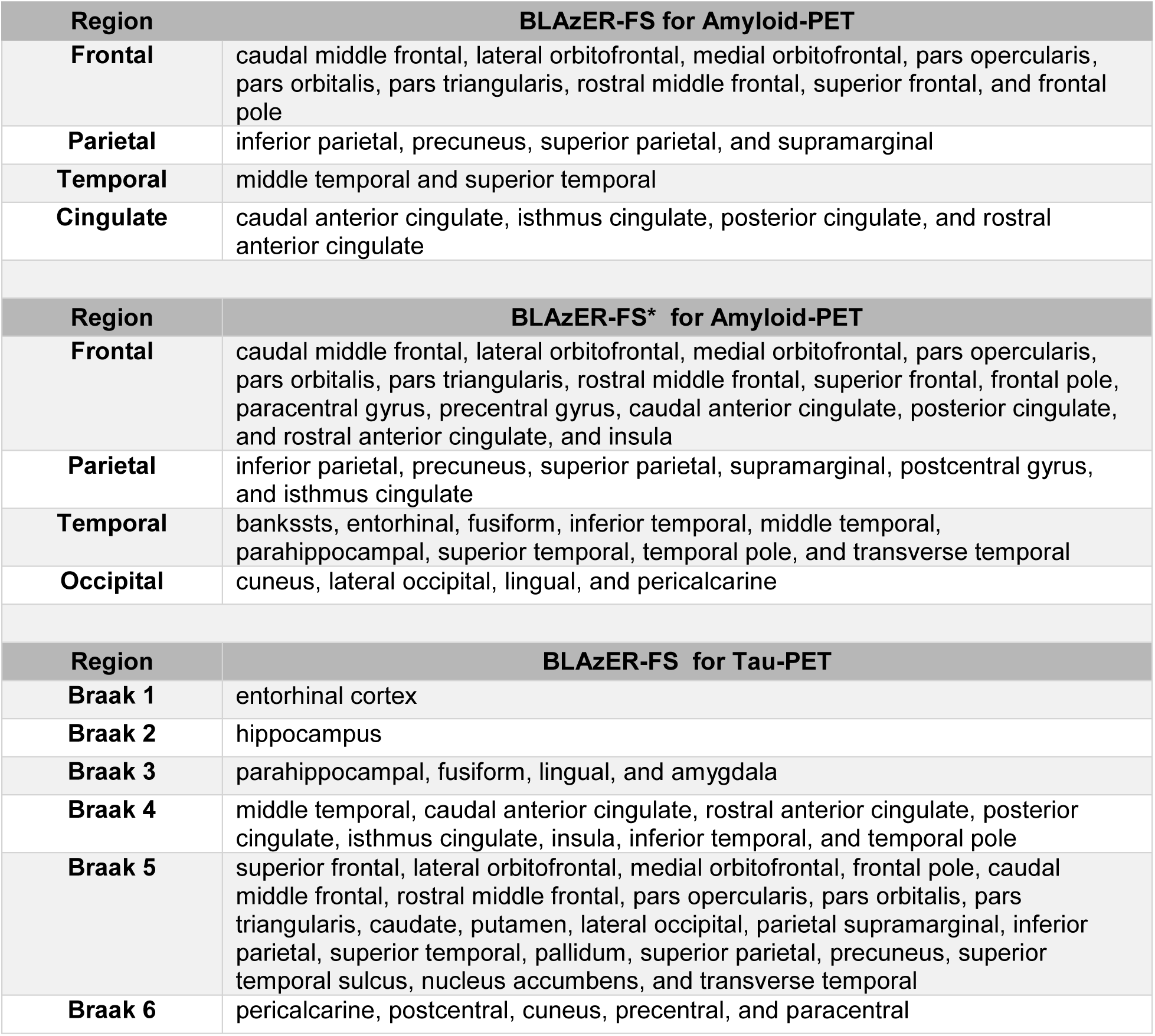
Anatomical Definitions of Brain Regions. Table of FreeSurfer (FS) subregions that comprised each of the cortices for amyloid-PET and each of the Braak staging regions for tau-PET. BLAzER-FS delineation was based on ADNI-defined brain subregions. However, for comparisons between BLAzER-FS and BLAzER-NR, BLAzER-FS anatomical regions were modified to match the NR-based regions of entire cortex. We denote this distinction as BLAzER-FS*.

| Region | BLAzER-FS for Amyloid-PET |
| --- | --- |
| **Frontal** | caudal middle frontal, lateral orbitofrontal, medial orbitofrontal, pars opercularis, pars orbitalis, pars triangularis, rostral middle frontal, superior frontal, and frontal pole |
| **Parietal** | inferior parietal, precuneus, superior parietal, and supramarginal |
| **Temporal** | middle temporal and superior temporal |
| **Cingulate** | caudal anterior cingulate, isthmus cingulate, posterior cingulate, and rostral anterior cingulate |
| Region | BLAzER-FS* for Amyloid-PET |
| **Frontal** | caudal middle frontal, lateral orbitofrontal, medial orbitofrontal, pars opercularis, pars orbitalis, pars triangularis, rostral middle frontal, superior frontal, frontal pole, paracentral gyrus, precentral gyrus, caudal anterior cingulate, posterior cingulate, and rostral anterior cingulate, and insula |
| **Parietal** | inferior parietal, precuneus, superior parietal, supramarginal, postcentral gyrus, and isthmus cingulate |
| **Temporal** | bankssts, entorhinal, fusiform, inferior temporal, middle temporal, parahippocampal, superior temporal, temporal pole, and transverse temporal |
| **Occipital** | cuneus, lateral occipital, lingual, and pericalcarine |
| Region | BLAzER-FS for Tau-PET |
| **Braak 1** | entorhinal cortex |
| **Braak 2** | hippocampus |
| **Braak 3** | parahippocampal, fusiform, lingual, and amygdala |
| **Braak 4** | middle temporal, caudal anterior cingulate, rostral anterior cingulate, posterior cingulate, isthmus cingulate, insula, inferior temporal, and temporal pole |
| **Braak 5** | superior frontal, lateral orbitofrontal, medial orbitofrontal, frontal pole, caudal middle frontal, rostral middle frontal, pars opercularis, pars orbitalis, pars triangularis, caudate, putamen, lateral occipital, parietal supramarginal, inferior parietal, superior temporal, pallidum, superior parietal, precuneus, superior temporal sulcus, nucleus accumbens, and transverse temporal |
| **Braak 6** | pericalcarine, postcentral, cuneus, precentral, and paracentral |

**Table S2.** ADNI vs. BLAzER: Volumetric Comparison. Regional comparison tables results for BLAzER vs. ADNI for volumetric MRI, showing SUVRs (mean ± standard deviation) with Pearson correlation coefficient across cognitively normal (CN), mild cognitively impaired (MCI), and Alzheimer’s dementia (AD) individuals across various brain regions. *** Denotes statistical significance of Pearson correlation p<0.0001.

| Volumetric MR |  |  |  |  |
| --- | --- | --- | --- | --- |
| Cognitive status | Brain regions | ADNI | BLAzER | r |
| CN | Frontal | 119.9 ± 13.5 | 116.2 ± 12.8 | 0.9547*** |
|  | Cingulate | 17.2 ± 2.3 | 17.2 ± 2.5 | 0.9548*** |
|  | Parietal | 74.3 ± 9.3 | 76.9 ± 8.6 | 0.9205*** |
|  | Temporal | 38.6 ± 5.5 | 40.7 ± 5.8 | 0.9623*** |
|  | Global | 249.9 ± 29.0 | 251.0 ± 27.5 | 0.9641*** |
| MCI | Frontal | 116.0 ± 17.0 | 112.9 ± 16.3 | 0.9761*** |
|  | Cingulate | 16.6 ± 2.8 | 16.4 ± 2.8 | 0.9570*** |
|  | Parietal | 71.6 ± 14.2 | 74.4 ± 14.2 | 0.9818*** |
|  | Temporal | 36.0 ± 7.0 | 38.2 ± 7.1 | 0.9702*** |
|  | Global | 240.2 ± 39.6 | 241.9 ± 38.6 | 0.9834*** |
| AD | Frontal | 122.8 ± 14.5 | 120.3 ± 14.4 | 0.9619*** |
|  | Cingulate | 17.7 ± 2.5 | 17.5 ± 2.8 | 0.9444*** |
|  | Parietal | 75.3 ± 11.1 | 79.4 ± 11.9 | 0.9536*** |
|  | Temporal | 37.6 ± 5.1 | 40.2 ± 5.5 | 0.9673*** |
|  | Global | 253.4 ± 30.8 | 257.4 ± 31.6 | 0.9685*** |

**Table S3.** ADNI vs. BLAzER: SUVR Comparison. Regional comparison tables results for BLAzER vs. ADNI for amyloid-PET (top) and tau-PET (bottom), showing SUVRs (mean ± standard deviation) with Pearson correlation coefficient across cognitively normal (CN), mild cognitively impaired (MCI), and Alzheimer’s dementia (AD) individuals across various brain regions. *** Denotes statistical significance of Pearson correlation p<0.0001.

| Amyloid-PET |  |  |  |  |
| --- | --- | --- | --- | --- |
| Cognitive status | Brain regions | ADNI | BLAzER | r |
| *CN* |  |  |  |  |
|  | Frontal | 1.11 ± 0.13 | 1.14 ± 0.13 | 0.9632*** |
|  | Cingulate | 1.24 ± 0.14 | 1.26 ± 0.15 | 0.9526*** |
|  | Parietal | 1.13 ± 0.14 | 1.16 ± 0.15 | 0.9643*** |
|  | Temporal | 1.04 ± 0.10 | 1.07 ± 0.11 | 0.9382*** |
|  | Global | 1.23 ± 0.19 | 1.16 ± 0.13 | 0.9894*** |
| *MCI* |  |  |  |  |
|  | Frontal | 1.22 ± 0.17 | 1.24 ± 0.17 | 0.9886*** |
|  | Cingulate | 1.31 ± 0.15 | 1.33 ± 0.15 | 0.9891*** |
|  | Parietal | 1.21 ± 0.16 | 1.23 ± 0.16 | 0.9781*** |
|  | Temporal | 1.13 ± 0.13 | 1.15 ± 0.14 | 0.9793*** |
|  | Global | 1.23 ± 0.17 | 1.24 ± 0.15 | 0.9884*** |
| *AD* |  |  |  |  |
|  | Frontal | 1.30 ± 0.20 | 1.32 ± 0.20 | 0.9956*** |
|  | Cingulate | 1.39 ± 0.19 | 1.40 ± 0.19 | 0.9945*** |
|  | Parietal | 1.31 ± 0.20 | 1.33 ± 0.190 | 0.9873*** |
|  | Temporal | 1.21 ± 0.18 | 1.23 ± 0.18 | 0.9870*** |
|  | Global | 1.20 ± 0.14 | 1.32 ± 0.19 | 0.9935*** |
| Tau-PET |  |  |  |  |
| Cognitive status | Brain regions | ADNI | BLAzER | r |
| *CN* |  |  |  |  |
|  | Braak 1 | 1.16 ± 0.16 | 1.30 ± 0.36 | 0.9705*** |
|  | Braak 2 | 1.22 ± 0.11 | 1.32 ± 0.21 | 0.9676*** |
|  | Braak 3 | 1.14 ± 0.10 | 1.28 ± 0.35 | 0.9894*** |
|  | Braak 4 | 1.14 ± 0.10 | 1.30 ± 0.37 | 0.9802*** |
|  | Braak 5 | 1.06 ± 0.08 | 1.21 ± 0.27 | 0.9822*** |
|  | Braak 6 | 0.99 ± 0.08 | 1.06 ± 0.12 | 0.9782*** |
| *MCI* |  |  |  |  |
|  | Braak 1 | 1.21 ± 0.24 | 1.26 ± 0.21 | 0.9806*** |
|  | Braak 2 | 1.34 ± 0.19 | 1.32 ± 0.19 | 0.9770*** |
|  | Braak 3 | 1.19 ± 0.09 | 1.19 ± 0.09 | 0.9097*** |
|  | Braak 4 | 1.19 ± 0.10 | 1.19 ± 0.11 | 0.9398*** |
|  | Braak 5 | 1.11 ± 0.09 | 1.14 ± 0.11 | 0.9417*** |
|  | Braak 6 | 1.02 ± 0.06 | 1.04 ± 0.09 | 0.9667*** |
| *AD* |  |  |  |  |
|  | Braak 1 | 1.44 ± 0.28 | 1.15 ± 0.18 | 0.9757*** |
|  | Braak 2 | 1.36 ± 0.22 | 1.24 ± 0.14 | 0.9821*** |
|  | Braak 3 | 1.38 ± 0.36 | 1.16 ± 0.12 | 0.9964*** |
|  | Braak 4 | 1.41 ± 0.46 | 1.21 ± 0.13 | 0.9986*** |
|  | Braak 5 | 1.22 ± 0.31 | 1.15 ± 0.10 | 0.9969*** |
|  | Braak 6 | 1.04 ± 0.14 | 1.01 ± 0.09 | 0.9896*** |

**Table S4.**
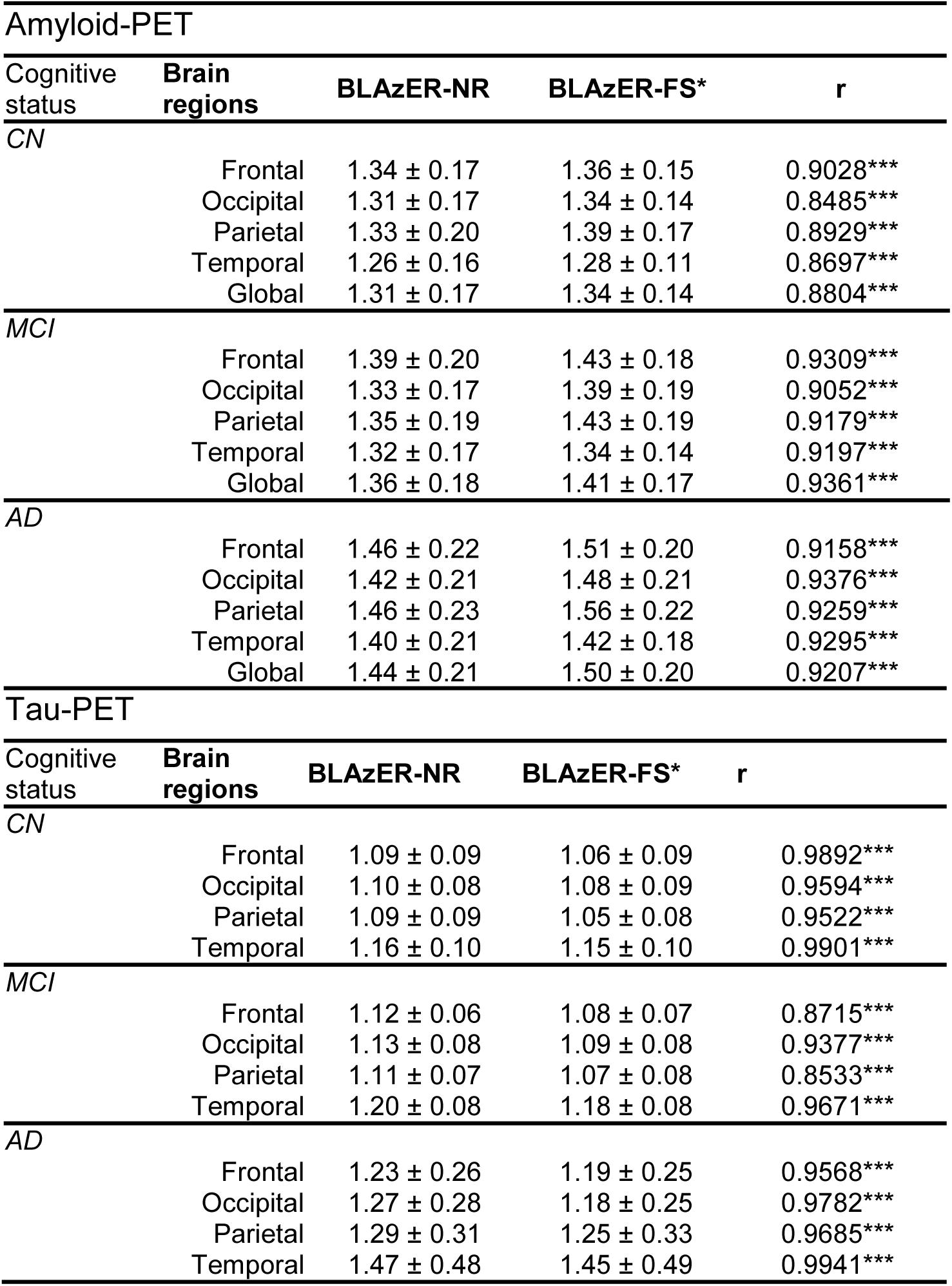
BLAzER-NR vs. BLAzER-FS*: SUVR Comparison. Regional comparison tables results for Neuroreader (BLAzER-NR) vs. FreeSurfer (BLAzER-FS*) utilizing BLAzER workflow for amyloid-PET (top) and tau-PET (bottom), showing SUVRs (mean ± standard deviation) with Pearson correlation coefficient across cognitively normal (CN), mild cognitively impaired (MCI), and Alzheimer’s dementia (AD) individuals across frontal, occipital, parietal, and temporal cortices as well as weighted global cortical average across these 4 brain regions. *Denotes BLAzER-FS utilizing NR regions. ***Denotes statistical significance of Pearson correlation p<0.0001.

| Amyloid-PET |  |  |  |  |
| --- | --- | --- | --- | --- |
| Cognitive status | Brain regions | BLAzER-NR | BLAzER-FS* | r |
| CN | Frontal | 1.34 ± 0.17 | 1.36 ± 0.15 | 0.9028*** |
|  | Occipital | 1.31 ± 0.17 | 1.34 ± 0.14 | 0.8485*** |
|  | Parietal | 1.33 ± 0.20 | 1.39 ± 0.17 | 0.8929*** |
|  | Temporal | 1.26 ± 0.16 | 1.28 ± 0.11 | 0.8697*** |
|  | Global | 1.31 ± 0.17 | 1.34 ± 0.14 | 0.8804*** |
| MCI | Frontal | 1.39 ± 0.20 | 1.43 ± 0.18 | 0.9309*** |
|  | Occipital | 1.33 ± 0.17 | 1.39 ± 0.19 | 0.9052*** |
|  | Parietal | 1.35 ± 0.19 | 1.43 ± 0.19 | 0.9179*** |
|  | Temporal | 1.32 ± 0.17 | 1.34 ± 0.14 | 0.9197*** |
|  | Global | 1.36 ± 0.18 | 1.41 ± 0.17 | 0.9361*** |
| AD | Frontal | 1.46 ± 0.22 | 1.51 ± 0.20 | 0.9158*** |
|  | Occipital | 1.42 ± 0.21 | 1.48 ± 0.21 | 0.9376*** |
|  | Parietal | 1.46 ± 0.23 | 1.56 ± 0.22 | 0.9259*** |
|  | Temporal | 1.40 ± 0.21 | 1.42 ± 0.18 | 0.9295*** |
|  | Global | 1.44 ± 0.21 | 1.50 ± 0.20 | 0.9207*** |
| Tau-PET |  |  |  |  |
| Cognitive status | Brain regions | BLAzER-NR | BLAzER-FS* | r |
| CN | Frontal | 1.09 ± 0.09 | 1.06 ± 0.09 | 0.9892*** |
|  | Occipital | 1.10 ± 0.08 | 1.08 ± 0.09 | 0.9594*** |
|  | Parietal | 1.09 ± 0.09 | 1.05 ± 0.08 | 0.9522*** |
|  | Temporal | 1.16 ± 0.10 | 1.15 ± 0.10 | 0.9901*** |
| MCI | Frontal | 1.12 ± 0.06 | 1.08 ± 0.07 | 0.8715*** |
|  | Occipital | 1.13 ± 0.08 | 1.09 ± 0.08 | 0.9377*** |
|  | Parietal | 1.11 ± 0.07 | 1.07 ± 0.08 | 0.8533*** |
|  | Temporal | 1.20 ± 0.08 | 1.18 ± 0.08 | 0.9671*** |
| AD | Frontal | 1.23 ± 0.26 | 1.19 ± 0.25 | 0.9568*** |
|  | Occipital | 1.27 ± 0.28 | 1.18 ± 0.25 | 0.9782*** |
|  | Parietal | 1.29 ± 0.31 | 1.25 ± 0.33 | 0.9685*** |
|  | Temporal | 1.47 ± 0.48 | 1.45 ± 0.49 | 0.9941*** |

## References

[1] Jack CR, Jr., Wiste HJ, Weigand SD, Therneau TM, Lowe VJ, Knopman DS, Gunter JL, Senjem ML, Jones DT, Kantarci K, Machulda MM, Mielke MM, Roberts RO, Vemuri P, Reyes DA, Petersen RC (2017) Defining imaging biomarker cut points for brain aging and Alzheimer’s disease. Alzheimers Dement 13, 205–216.

[2] Hutton C, Declerck J, Mintun MA, Pontecorvo MJ, Devous MD Sr., Joshi AD, Alzheimer’s Disease Neuroimaging I (2015) Quantification of 18F-florbetapir PET: comparison of two analysis methods. Eur J Nucl Med Mol Imaging 42, 725–732.

[3] Bullich S, Seibyl J, Catafau AM, Jovalekic A, Koglin N, Barthel H, Sabri O, De Santi S (2017) Optimized classification of (18)F-Florbetaben PET scans as positive and negative using an SUVR quantitative approach and comparison to visual assessment. Neuroimage Clin 15, 325–332.

[4] Minoshima S, Koeppe RA, Frey KA, Ishihara M, Kuhl DE (1994) Stereotactic PET atlas of the human brain: aid for visual interpretation of functional brain images. J Nucl Med 35, 949–954.

[5] Destrieux C, Fischl B, Dale A, Halgren E (2010) Automatic parcellation of human cortical gyri and sulci using standard anatomical nomenclature. Neuroimage 53, 1–15.

[6] Lehmann M, Douiri A, Kim LG, Modat M, Chan D, Ourselin S, Barnes J, Fox NC (2010) Atrophy patterns in Alzheimer’s disease and semantic dementia: a comparison of FreeSurfer and manual volumetric measurements. Neuroimage 49, 2264–2274.

[7] Fischl B (2012) FreeSurfer. Neuroimage 62, 774–781.

[8] Fischl B, Salat DH, Busa E, Albert M, Dieterich M, Haselgrove C, van der Kouwe A, Killiany R, Kennedy D, Klaveness S, Montillo A, Makris N, Rosen B, Dale AM (2002) Whole brain segmentation: automated labeling of neuroanatomical structures in the human brain. Neuron 33, 341–355.

[9] Fischl B, Salat DH, van der Kouwe AJ, Makris N, Segonne F, Quinn BT, Dale AM (2004) Sequence-independent segmentation of magnetic resonance images. Neuroimage 23 Suppl 1, S69–84.

[10] Ahdidan J, Raji CA, DeYoe EA, Mathis J, Noe KO, Rimestad J, Kjeldsen TK, Mosegaard J, Becker JT, Lopez O (2016) Quantitative Neuroimaging Software for Clinical Assessment of Hippocampal Volumes on MR Imaging. J Alzheimers Dis 49, 723–732.

[11] Tanpitukpongse TP, Mazurowski MA, Ikhena J, Petrella JR, Alzheimer’s Disease Neuroimaging I (2017) Predictive Utility of Marketed Volumetric Software Tools in Subjects at Risk for Alzheimer Disease: Do Regions Outside the Hippocampus Matter? AJNR Am J Neuroradiol 38, 546–552.

[12] Bredesen DE, Amos EC, Canick J, Ackerley M, Raji C, Fiala M, Ahdidan J (2016) Reversal of cognitive decline in Alzheimer’s disease. Aging (Albany NY) 8, 1250–1258.

[13] Chen BT, Sethi SK, Jin T, Patel SK, Ye N, Sun CL, Rockne RC, Haacke EM, Root JC, Saykin AJ, Ahles TA, Holodny AI, Prakash N, Mortimer J, Waisman J, Yuan Y, Somlo G, Li D, Yang R, Tan H, Katheria V, Morrison R, Hurria A (2018) Assessing brain volume changes in older women with breast cancer receiving adjuvant chemotherapy: a brain magnetic resonance imaging pilot study. Breast Cancer Res 20, 38.

[14] Schain M, Varnas K, Cselenyi Z, Halldin C, Farde L, Varrone A (2014) Evaluation of two automated methods for PET region of interest analysis. Neuroinformatics 12, 551–562.

[15] Lundqvist R, Lilja J, Thomas BA, Lotjonen J, Villemagne VL, Rowe CC, Thurfjell L (2013) Implementation and validation of an adaptive template registration method for 18F-flutemetamol imaging data. J Nucl Med 54, 1472–1478.

[16] Gorgolewski K, Burns CD, Madison C, Clark D, Halchenko YO, Waskom ML, Ghosh SS (2011) Nipype: a flexible, lightweight and extensible neuroimaging data processing framework in python. Front Neuroinform 5, 13.

[17] Su Y, D’Angelo GM, Vlassenko AG, Zhou G, Snyder AZ, Marcus DS, Blazey TM, Christensen JJ, Vora S, Morris JC, Mintun MA, Benzinger TL (2013) Quantitative analysis of PiB-PET with FreeSurfer ROIs. PLoS One 8, e73377.

[18] Cusack R, Vicente-Grabovetsky A, Mitchell DJ, Wild CJ, Auer T, Linke AC, Peelle JE (2014) Automatic analysis (aa): efficient neuroimaging workflows and parallel processing using Matlab and XML. Front Neuroinform 8, 90.

[19] Tzourio-Mazoyer N, Landeau B, Papathanassiou D, Crivello F, Etard O, Delcroix N, Mazoyer B, Joliot M (2002) Automated anatomical labeling of activations in SPM using a macroscopic anatomical parcellation of the MNI MRI single-subject brain. Neuroimage 15, 273–289.

[20] Frings L, Hellwig S, Bormann T, Spehl TS, Buchert R, Meyer PT (2018) Amyloid load but not regional glucose metabolism predicts conversion to Alzheimer’s dementia in a memory clinic population. Eur J Nucl Med Mol Imaging.

[21] Landau SM, Fero A, Baker SL, Koeppe R, Mintun M, Chen K, Reiman EM, Jagust WJ (2015) Measurement of longitudinal beta-amyloid change with 18F-florbetapir PET and standardized uptake value ratios. J Nucl Med 56, 567–574.

[22] Martinez G, Vernooij RW, Fuentes Padilla P, Zamora J, Bonfill Cosp X, Flicker L (2017) 18F PET with florbetapir for the early diagnosis of Alzheimer’s disease dementia and other dementias in people with mild cognitive impairment (MCI). Cochrane Database Syst Rev 11, CD012216.

[23] Martinez G, Vernooij RW, Fuentes Padilla P, Zamora J, Flicker L, Bonfill Cosp X (2017) 18F PET with florbetaben for the early diagnosis of Alzheimer’s disease dementia and other dementias in people with mild cognitive impairment (MCI). Cochrane Database Syst Rev 11, CD012883.

[24] Martinez G, Vernooij RW, Fuentes Padilla P, Zamora J, Flicker L, Bonfill Cosp X (2017) 18F PET with flutemetamol for the early diagnosis of Alzheimer’s disease dementia and other dementias in people with mild cognitive impairment (MCI). Cochrane Database Syst Rev 11, CD012884.

[25] Cummings J (2018) The National Institute on Aging-Alzheimer’s Association Framework on Alzheimer’s disease: Application to clinical trials. Alzheimers Dement.

[26] Jack CR,Jr., Bennett DA, Blennow K, Carrillo MC, Dunn B, Haeberlein SB, Holtzman DM, Jagust W, Jessen F, Karlawish J, Liu E, Molinuevo JL, Montine T, Phelps C, Rankin KP, Rowe CC, Scheltens P, Siemers E, Snyder HM, Sperling R, Contributors (2018) NIA-AA Research Framework: Toward a biological definition of Alzheimer’s disease. Alzheimers Dement 14, 535–562.

[27] Jack CR, Jr., Wiste HJ, Weigand SD, Therneau TM, Knopman DS, Lowe V, Vemuri P, Mielke MM, Roberts RO, Machulda MM, Senjem ML, Gunter JL, Rocca WA, Petersen RC (2017) Age-specific and sex-specific prevalence of cerebral beta-amyloidosis, tauopathy, and neurodegeneration in cognitively unimpaired individuals aged 50-95 years: a cross-sectional study. Lancet Neurol 16, 435–444.

[28] Vemuri P, Lowe VJ, Knopman DS, Senjem ML, Kemp BJ, Schwarz CG, Przybelski SA, Machulda MM, Petersen RC, Jack CR, Jr. (2017) Tau-PET uptake: Regional variation in average SUVR and impact of amyloid deposition. Alzheimers Dement (Amst) 6, 21–30.

[29] Friedland RP, Budinger TF, Ganz E, Yano Y, Mathis CA, Koss B, Ober BA, Huesman RH, Derenzo SE (1983) Regional cerebral metabolic alterations in dementia of the Alzheimer type: positron emission tomography with [18F]fluorodeoxyglucose. J Comput Assist Tomogr 7, 590–598.

[30] Mosconi L, Tsui WH, De Santi S, Li J, Rusinek H, Convit A, Li Y, Boppana M, de Leon MJ (2005) Reduced hippocampal metabolism in MCI and AD: automated FDG-PET image analysis. Neurology 64, 1860–1867.

[31] Edison P, Carter SF, Rinne JO, Gelosa G, Herholz K, Nordberg A, Brooks DJ, Hinz R (2013) Comparison of MRI based and PET template based approaches in the quantitative analysis of amyloid imaging with PIB-PET. Neuroimage 70, 423–433.

[32] Evans AC, Marrett S, Neelin P, Collins L, Worsley K, Dai W, Milot S, Meyer E, Bub D (1992) Anatomical mapping of functional activation in stereotactic coordinate space. Neuroimage 1, 43–53.

[33] Landau SM, Breault C, Joshi AD, Pontecorvo M, Mathis CA, Jagust WJ, Mintun MA, Alzheimer’s Disease Neuroimaging I (2013) Amyloid-beta imaging with Pittsburgh compound B and florbetapir: comparing radiotracers and quantification methods. J Nucl Med 54, 70–77.

[34] Landau SM, Mintun MA, Joshi AD, Koeppe RA, Petersen RC, Aisen PS, Weiner MW, Jagust WJ, Alzheimer’s Disease Neuroimaging I (2012) Amyloid deposition, hypometabolism, and longitudinal cognitive decline. Ann Neurol 72, 578–586.

[35] Aalto S, Scheinin NM, Kemppainen NM, Nagren K, Kailajarvi M, Leinonen M, Scheinin M, Rinne JO (2009) Reproducibility of automated simplified voxel-based analysis of PET amyloid ligand [11C]PIB uptake using 30-min scanning data. Eur J Nucl Med Mol Imaging 36, 1651–1660.

[36] Jagust WJ, Landau SM, Koeppe RA, Reiman EM, Chen K, Mathis CA, Price JC, Foster NL, Wang AY (2015) The Alzheimer’s Disease Neuroimaging Initiative 2 PET Core: 2015. Alzheimers Dement 11, 757–771.

[37] Scholl M, Lockhart SN, Schonhaut DR, O’Neil JP, Janabi M, Ossenkoppele R, Baker SL, Vogel JW, Faria J, Schwimmer HD, Rabinovici GD, Jagust WJ (2016) PET Imaging of Tau Deposition in the Aging Human Brain. Neuron 89, 971–982.

[38] Eisenstein SA, Koller JM, Piccirillo M, Kim A, Antenor-Dorsey JA, Videen TO, Snyder AZ, Karimi M, Moerlein SM, Black KJ, Perlmutter JS, Hershey T (2012) Characterization of extrastriatal D2 in vivo specific binding of [(1)(8)F](N-methyl)benperidol using PET. Synapse 66, 770–780.

[39] Savli M, Bauer A, Mitterhauser M, Ding YS, Hahn A, Kroll T, Neumeister A, Haeusler D, Ungersboeck J, Henry S, Isfahani SA, Rattay F, Wadsak W, Kasper S, Lanzenberger R (2012) Normative database of the serotonergic system in healthy subjects using multi-tracer PET. Neuroimage 63, 447–459.

[40] van Mourik T, Snoek L, Knapen T, Norris DG (2018) Porcupine: A visual pipeline tool for neuroimaging analysis. PLoS Comput Biol 14, e1006064.

[41] Braak H, Alafuzoff I, Arzberger T, Kretzschmar H, Del Tredici K (2006) Staging of Alzheimer disease-associated neurofibrillary pathology using paraffin sections and immunocytochemistry. Acta Neuropathol 112, 389–404.

[42] Clark CM, Schneider JA, Bedell BJ, Beach TG, Bilker WB, Mintun MA, Pontecorvo MJ, Hefti F, Carpenter AP, Flitter ML, Krautkramer MJ, Kung HF, Coleman RE, Doraiswamy PM, Fleisher AS, Sabbagh MN, Sadowsky CH, Reiman EP, Zehntner SP, Skovronsky DM, Group AAS (2011) Use of florbetapir-PET for imaging beta-amyloid pathology. JAMA 305, 275–283.

[43] Yang J, Hu C, Guo N, Dutta J, Vaina LM, Johnson KA, Sepulcre J, Fakhri GE, Li Q (2017) Partial volume correction for PET quantification and its impact on brain network in Alzheimer’s disease. Sci Rep 7, 13035.

[44] Passamonti L, Vazquez Rodriguez P, Hong YT, Allinson KS, Williamson D, Borchert RJ, Sami S, Cope TE, Bevan-Jones WR, Jones PS, Arnold R, Surendranathan A, Mak E, Su L, Fryer TD, Aigbirhio FI, O’Brien JT, Rowe JB (2017) 18F-AV-1451 positron emission tomography in Alzheimer’s disease and progressive supranuclear palsy. Brain 140, 781–791.

[45] Choi JY, Cho H, Ahn SJ, Lee JH, Ryu YH, Lee MS, Lyoo CH (2018) Off-Target (18)F-AV-1451 Binding in the Basal Ganglia Correlates with Age-Related Iron Accumulation. J Nucl Med 59, 117–120.

[46] Gronenschild EH, Habets P, Jacobs HI, Mengelers R, Rozendaal N, van Os J, Marcelis M (2012) The effects of FreeSurfer version, workstation type, and Macintosh operating system version on anatomical volume and cortical thickness measurements. PLoS One 7, e38234.

[47] Saint-Aubert L, Nemmi F, Peran P, Barbeau EJ, Payoux P, Chollet F, Pariente J (2014) Comparison between PET template-based method and MRI-based method for cortical quantification of florbetapir (AV-45) uptake in vivo. Eur J Nucl Med Mol Imaging 41, 836–843.

[48] Tuszynski T, Rullmann M, Luthardt J, Butzke D, Tiepolt S, Gertz HJ, Hesse S, Seese A, Lobsien D, Sabri O, Barthel H (2016) Evaluation of software tools for automated identification of neuroanatomical structures in quantitative beta-amyloid PET imaging to diagnose Alzheimer’s disease. Eur J Nucl Med Mol Imaging 43, 1077–1087.

[49] Thurfjell L, Lilja J, Lundqvist R, Buckley C, Smith A, Vandenberghe R, Sherwin P (2014) Automated quantification of 18F-flutemetamol PET activity for categorizing scans as negative or positive for brain amyloid: concordance with visual image reads. J Nucl Med 55, 1623–1628.

[50] Dupont AC, Largeau B, Santiago Ribeiro MJ, Guilloteau D, Tronel C, Arlicot N (2017) Translocator Protein-18 kDa (TSPO) Positron Emission Tomography (PET) Imaging and Its Clinical Impact in Neurodegenerative Diseases. Int J Mol Sci 18.

[51] Largeau B, Dupont AC, Guilloteau D, Santiago-Ribeiro MJ, Arlicot N (2017) TSPO PET Imaging: From Microglial Activation to Peripheral Sterile Inflammatory Diseases? Contrast Media Mol Imaging 2017, 6592139.

[52] Selvaraj S, Bloomfield PS, Cao B, Veronese M, Turkheimer F, Howes OD (2018) Brain TSPO imaging and gray matter volume in schizophrenia patients and in people at ultra high risk of psychosis: An [(11)C]PBR28 study. Schizophr Res 195, 206–214.

[53] Lin KJ, Hsu WC, Hsiao IT, Wey SP, Jin LW, Skovronsky D, Wai YY, Chang HP, Lo CW, Yao CH, Yen TC, Kung MP (2010) Whole-body biodistribution and brain PET imaging with [18F]AV-45, a novel amyloid imaging agent--a pilot study. Nucl Med Biol 37, 497–508.

